# NFATc acts as a non-canonical phenotypic stability factor for a hybrid epithelial/mesenchymal phenotype

**DOI:** 10.1101/2020.04.18.047803

**Authors:** Ayalur Raghu Subbalakshmi, Deepali Kundnani, Kuheli Biswas, Anandamohan Ghosh, Samir M Hanash, Satyendra C Tripathi, Mohit Kumar Jolly

## Abstract

Metastasis remains the cause of over 90% of cancer-related deaths. Cells undergoing metastasis use phenotypic plasticity to adapt to their changing environmental conditions and avoid therapy and immune response. Reversible transitions between epithelial and mesenchymal phenotypes - Epithelial-Mesenchymal Transition (EMT) and its reverse Mesenchymal-Epithelial Transition (MET) - form a key axis of phenotypic plasticity during metastasis and therapy resistance. Recent studies have shown that the cells undergoing EMT/MET can attain one or more hybrid epithelial/mesenchymal (E/M) phenotypes, the process of which is termed as partial EMT/MET. Cells in hybrid E/M phenotype(s) can be more aggressive than those in either epithelial or mesenchymal state. Thus, it is crucial to identify the factors and regulatory networks enabling such hybrid E/M phenotypes. Here, employing an integrated computational-experimental approach, we show that the transcription factor NFATc can inhibit the process of complete EMT, thus stabilizing the hybrid E/M phenotype. It increases the range of parameters enabling the existence of a hybrid E/M phenotype, thus behaving as a phenotypic stability factor (PSF). However, unlike previously identified PSFs, it does not increase the mean residence time of the cells in hybrid E/M phenotypes, as shown by stochastic simulations; rather it enables the co-existence of epithelial, mesenchymal and hybrid E/M phenotypes and transitions among them. Clinical data suggests the effect of NFATc on patient survival in a tissue-specific or context-dependent manner. Together, our results indicate that NFATc behaves as a non-canonical phenotypic stability factor for a hybrid E/M phenotype.

## Introduction

Metastasis remains clinically insuperable and causes over 90% of cancer related deaths (1). A hallmark of metastasizing cells is phenotypic plasticity, which empowers them to adapt to their ever-changing microenvironment, while evading therapy and immune response (2). Cells displaying phenotypic plasticity can have profound consequences: an identical genetic background can give rise to varying phenotypes under different environmental conditions, enabling non-genetic heterogeneity (3,4). A crucial axis of phenotypic plasticity during metastasis is epithelial-mesenchymal plasticity, which allows bidirectional switching of cells among an epithelial phenotype, a mesenchymal phenotype, and one or more hybrid epithelial/mesenchymal (E/M) phenotypes (5). These hybrid E/M cells can be more metastatic than cells in epithelial or mesenchymal states (6,7) and can exhibit collective cell migration as clusters of circulating tumour cells (CTCs) (8–10) – the major drivers of metastasis (11). Thus, understanding the molecular mechanisms enabling one or more hybrid E/M phenotype(s) is key to decoding and eventually restricting metastasis.

EMT is influenced by various pathways such as transforming growth factor β (TGF-β), Wnt–β-catenin, bone morphogenetic protein (BMP), Notch, Hedgehog, and receptor tyrosine kinases (12). These EMT signals alter the levels of one or more EMT-inducing transcription factors (EMT-TFs) such as ZEB and SNAIL which can directly repress various epithelial molecules such as E-cadherin and/or induce the expression of various mesenchymal ones (5). ZEB and SNAIL form mutually inhibitory feedback loops with two microRNA families miR-200 and miR-34 where the transcription factors and the micro-RNAs mutually inhibit each other (13–16). Overexpression of ZEB promotes EMT and silences the micro-RNAs which act as a safeguard for maintaining an epithelial phenotype (13). Recent studies have indicated the involvement of phenotypic stability factors (PSF) such as GRHL2, OVOL2, NUMB and NRF2 that can maintain the cells in a hybrid E/M phenotype(s) and prevent the cells from undergoing a complete EMT (17–22). Knockdown of these PSFs usually drove hybrid E/M cells towards a completely mesenchymal phenotype, as observed in H1975 non-small cell lung cancer (NSCLC) cells which can maintain a hybrid E/M phenotype stably over multiple passages *in vitro* (20). Higher levels of these PSFs also increased the mean residence times of cells in a hybrid E/M phenotype (23) and associated with poor patient survival, thus highlighting the clinical significance of hybrid E/M phenotypes (24).

Here, we investigate the role of the Nuclear factor of activated T-cell (NFATc) in mediating EMT. NFATc is a family of five transcription factors (NFAT1-5), four of which (NFATc1-4) are regulated by calcium Ca^2+^ signalling (25). Initially identified as functionally important for T lymphocytes, the NFAT family regulates cell cycle progression, gene expression and apoptosis (25). Abnormalities in NFATc signaling have been reported in many carcinomas as well as lymphoma and leukemia (26). Recent evidence has suggested the interconnections of NFATc with EMT circuitry. On one hand, overexpression of NFATc increased the levels of TWIST, ZEB1, SNAI1; its downregulation decreased the levels of these EMT-TFs as well as mesenchymal markers such as N-cadherin and Vimentin ZEB (27). On the other hand, NFATc transcriptional activity was also shown to be crucial for maintaining E-cadherin levels (28), which can inhibit ZEB1 indirectly (29,30) and restrict EMT. Moreover, NFATc can activate SOX2 (27,31) which can upregulate levels of miR-200 (32); overexpression of miR-200 can drive MET (33). These opposing interactions of NFATc with EMT circuitry lead to the question of whether NFATc promotes or inhibit EMT. Here, we designed a mechanism-based mathematical model that captures the interconnections between NFATc signaling and EMT circuitry. Our analysis predicts that NFATc can stabilize a hybrid E/M phenotype, facilitating cellular plasticity. Knockdown of NFATc in H1975 cells pushed the hybrid E/M cells into a mesenchymal state, validating our prediction that NFATc can function as a PSF.

## Results

### NFATc inhibits the progression of complete EMT

To determine the role of NFATc in EMT, we first investigated the dynamics of crosstalk between NFATc and an EMT regulatory circuit (shown in the dotted rectangle) that includes miR-200, ZEB and SNAIL (Fig 1A). This crosstalk was captured through a set of coupled ordinary differential equations (ODEs).

**Fig1:**
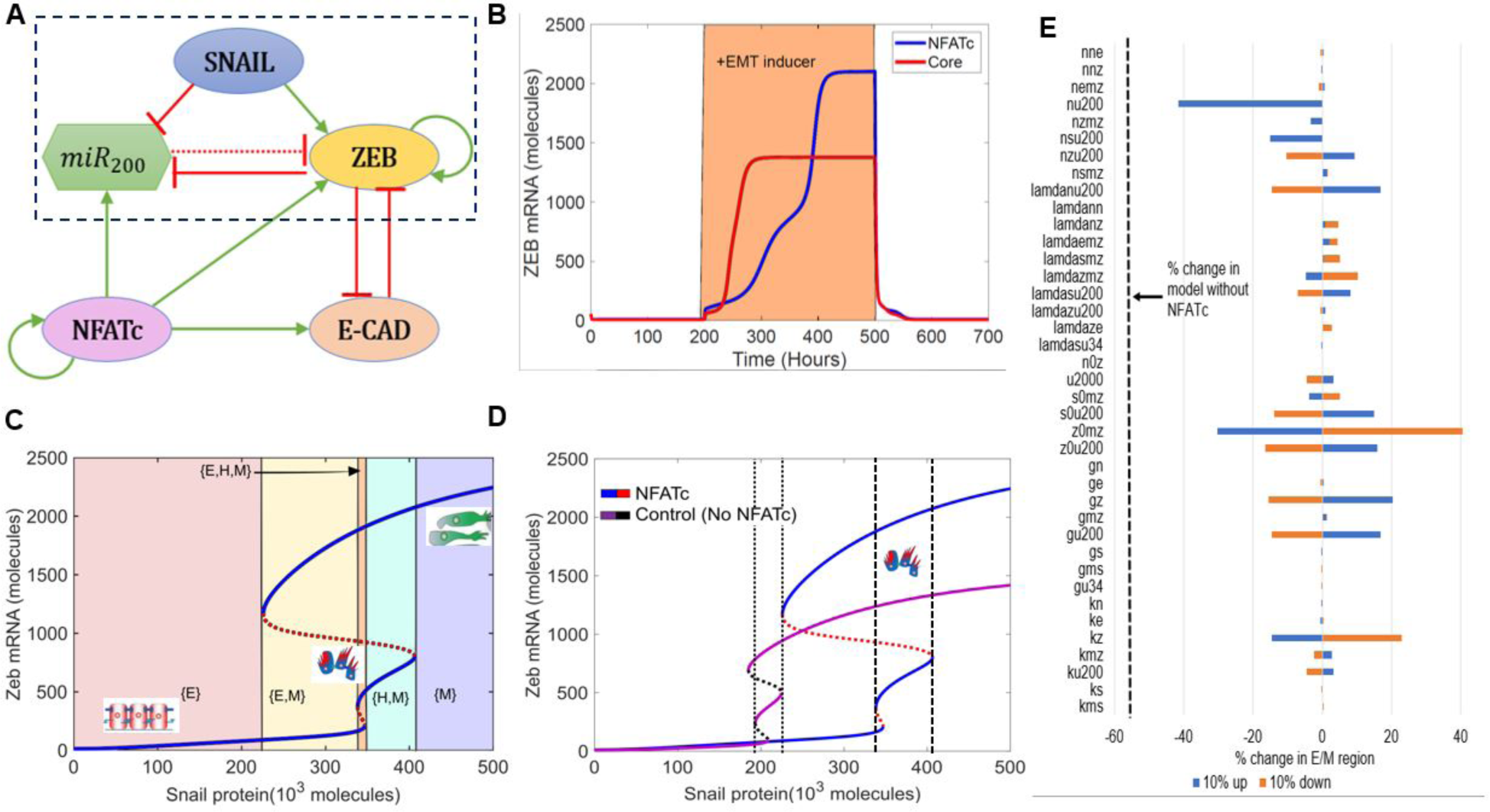
NFATc inhibits the progression of complete EMT. A) Schematic representation of the EMT network coupled with NFATc. Red bars denote inhibition, green arrows indicate activation. Solid arrows represent transcriptional regulation and dotted line represent micro-RNA mediated regulation. B) Temporal dynamics of ZEB mRNA levels in a cell starting in an epithelial phenotype, when exposed to a high level of S=330000 molecules (orange-shaded region) for circuit shown in A (NFATc) and that shown in the dotted rectangle in A (core). C) Bifurcation diagram of ZEB mRNA levels as driven by SNAIL signal for the circuit shown in A. Curves denote the value of ZEB mRNA upon equilibration, where continuous curves stand for stable steady states and dotted lines represent unstable solutions. Different coloured shaded regions show the existence/co-existence of different phenotypes; cartoons depict epithelial (E), hybrid E/M (H) and mesenchymal (M) states. D) Bifurcation diagrams indicating the ZEB mRNA levels for increasing SNAIL levels for the coupled EMT-NFATc circuit (solid blue and dotted red curve) and the core EMT circuit (solid purple and dotted black curve). E) Sensitivity analysis indicating percent change in the interval of SNAIL levels for stable hybrid E/M region, when corresponding parameter values are varied by ± 10%. The black dotted line indicates the percent change in the stable hybrid region in the absence of NFATc (core network) when compared to the coupled network with NFATc.

First, we examined the temporal dynamics of a cell in response to SNAIL levels. SNAIL represents the effect of an exogenous EMT-inducing signal such as TGF-β signaling. In the absence of NFATc, a cell that started in an epithelial state (high miR-200, low ZEB) first transitioned to a hybrid E/M phenotype and later to a mesenchymal state (low miR-200, high ZEB). The presence of NFATc, however, delayed this transition to a mesenchymal state (Fig 1B, S1A). Interestingly, the steady state value of ZEB mRNA levels was higher in case of NFATc-EMT coupled network as compared to the control case (circuit bounded by the dotted rectangle in Fig 1A); this difference can be ascribed to the activation of ZEB by NFATc (Fig 1B). Indeed, strengthening the activation of miR-200 by NFATc and/or weakening the activation of ZEB by NFATc prevented upregulation of ZEB levels and consequent attainment of the mesenchymal phenotype (Fig S1B,C).

Next, we calculated a bifurcation diagram of cellular EMT phenotypes in response to increasing levels of SNAIL, an external EMT-inducing signal (Fig 1B). We observe that the system switches from epithelial to hybrid E/M and then to mesenchymal state with increasing SNAIL signal, as indicated by increasing values of ZEB mRNA and decreasing values of miR-200 (Fig 1C, S1D). Next, we compared the bifurcation diagram of the NFATc coupled network with that of the control case (i.e. without NFATc –the circuit bounded by the dotted rectangles in Fig 1A) to determine the changes in the system behaviour conferred by NFATc (Fig 1D). In the presence of NFATc, a higher value of SNAIL, i.e. a stronger external signal, was required for the cells to exit the epithelial phenotype. Moreover, in the presence of NFATc, the cell maintained a hybrid E/M phenotype over a broader range of SNAIL levels (compare the range of SNAIL levels bounded by dotted lines vs. dashed lines in Fig 1D), thus requiring a much stronger stimulus to undergo a complete EMT.

Finally, to ascertain the robustness of the effect of NFATc in associating a larger range of SNAIL values for the existence of hybrid E/M phenotype, a sensitivity analysis was performed where each parameter of the model was varied – one at a time – by ± 10%, and the corresponding change in the range of values of SNAIL enabling a hybrid E/M phenotype (i.e. the interval of x-axis between dashed lines) was measured. For most of the model parameters, the relative change in this range of values was quite small, suggesting the robustness of the model predictions (Fig 1E). A change in few selected parameters such as the interaction between NFATc and miR200, and self-activation of ZEB exhibited stronger sensitivity; nonetheless, even in these few cases, the decrease in range of SNAIL levels enabling a hybrid E/M phenotype is smaller when compared to the case in absence of NFATc (dotted line in Fig1E). Put together, these observations suggest that NFATc may inhibit the progression to a complete EMT and can behave as a ‘phenotypic stability factor’ (PSF) for hybrid E/M phenotype.

### Knockdown of NFATc in H1975 cells promotes complete EMT

To validate our model prediction that NFATc functions as a PSF for the hybrid E/M phenotype, we knocked down NFATc1 using siRNAs in non-small cell lung cancer (NSCLC) H1975 cells with a stable hybrid E/M phenotype.

Individual H1975 cells can co-express E-cadherin and Vimentin (20). We observed that cells treated with NFATc1 siRNAs mostly lost E-cadherin staining (Fig 2A). NFATc knockdown decreased the E-cadherin levels and increased the levels of ZEB, SNAIL and Vimentin, both at protein and mRNA levels (FigS2A-C). Thus, NFATc1 knockdown can push stable hybrid E/M cells into a mesenchymal state. Additionally, NFATc1 knockdown reduced the migration potential of H1975 cells as observed in scratch and trans-well migration assays. These findings indicate that the hybrid E/M cells may exhibit greater migratory and invasive potential when compared to mesenchymal cells (Fig 2B-C), reminiscent of our previous observations comparing hybrid E/M cells with mesenchymal ones (17). Overall, these experimental results provide a proof-of-principle validation of our model predictions that NFATc can stabilize a hybrid E/M phenotype.

**Fig2:**
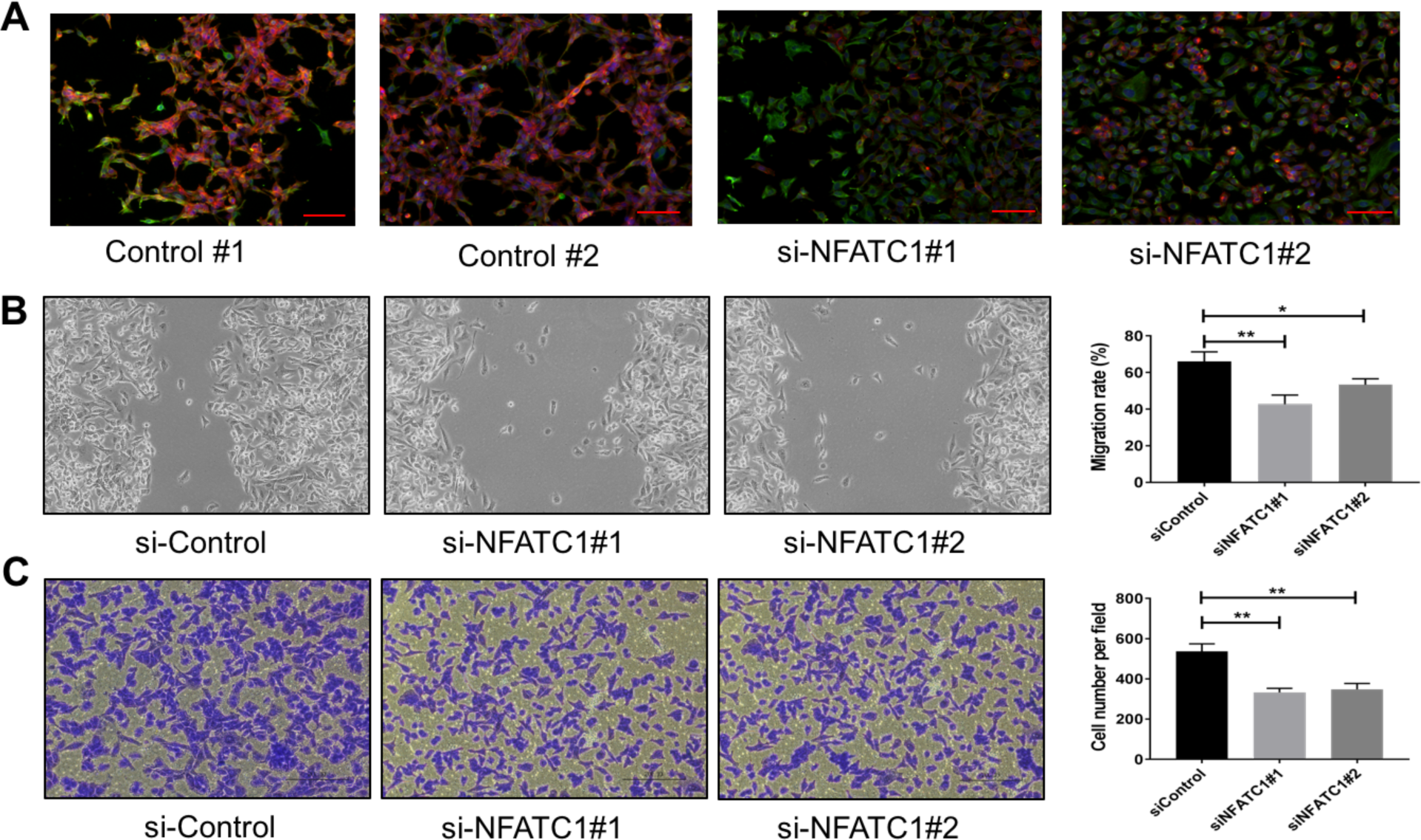
NFATc knockdown in H1975 cells promotes progression towards complete EMT. A) Expression of CDH1 (E-cadherin, Red) and VIM (Vimentin, Green) examined by immuno-fluorescence staining in H1975 cells for control and NFATc1 knockdown case. Scale bar 100 µm. B) Scratch assay for control H1975 cells and those treated with siRNAs against NFATc. Magnification:100X (quantification in last column). C) Same as B but for trans-well migration assay. *p < 0.05, **p<0.005, ***p<0.001 using Students’ t-test; n=3.

### NFATc does not increase the mean residence time of the hybrid E/M phenotype

In addition to extending the range of SNAIL levels enabling a hybrid E/M phenotype, the previously identified PSFs – GRHL2, OVOL1/2, and ΔNP63α – had another trait: their presence increased the mean residence time (MRT) of cells in hybrid E/M phenotype. MRT is the average time spent by the cells in a particular phenotype (basin of attraction) – E, M and hybrid E/M – calculated via stochastic simulations (23). Thus, the phenotype with a larger MRT implies a relatively higher stability of the same, as compared to other co-existing phenotypes/states. Hence, beyond enabling a larger range of values of SNAIL (or any other EMT inducing signal) for the existence of a hybrid E/M phenotype (as shown via bifurcation diagrams), increased MRT can be considered as another hallmark trait of a PSF. We next investigated whether NFATc increased the MRT of cells in a hybrid E/M phenotype.

Even though NFATc extended the range of SNAIL values enabling a hybrid E/M state, the hybrid E/M state always co-existed with epithelial and/or mesenchymal states ({E, H, M} and {H, M} phases in Fig 1B); no monostable regime ({H}) for a hybrid E/M state was seen in the case of NFATc, as observed with GRHL2, OVOL1/2, ΔNP63α, NUMB and NRF2 (17–21,34). Similarly, compared to the other PSFs, the presence of NFATc does not increase the absolute value of MRT for a hybrid E/M phenotype as compared to the case without NFATc. In the case of control circuit, the MRT of the epithelial state is higher than that of the mesenchymal state in the {E, M} bi-stable phase. In the {H, M} phase, the MRT of the mesenchymal state dominates that of the hybrid E/M state as SNAIL values are increased (Fig 3A). Similar trends are seen in the case of NFATc-EMT coupled network; however, in the case of {E, H, M} phase in presence of NFATc, the MRT of hybrid E/M state is not higher as compared to that of epithelial or mesenchymal states (compare the red curve in Fig 3B vs. that in Fig 3A). This trend is also seen in the barrier height calculated from the potential difference between the local minimum and saddle points corresponding to these states (Fig. 3C,D). We also plotted the potential landscapes for the NFATc-EMT coupled network at varying SNAIL levels (Fig S3), which were consistent with the trend of barrier heights seen; for instance, at S=323*10^3^ or S=330*10^3^ molecules, the barrier height of hybrid E/M state was more than that of mesenchymal state, but at S=380*10^3^, that of mesenchymal state was higher. Put together, these results suggest NFATc does not increase the MRT of hybrid E/M state.

**Fig3:**
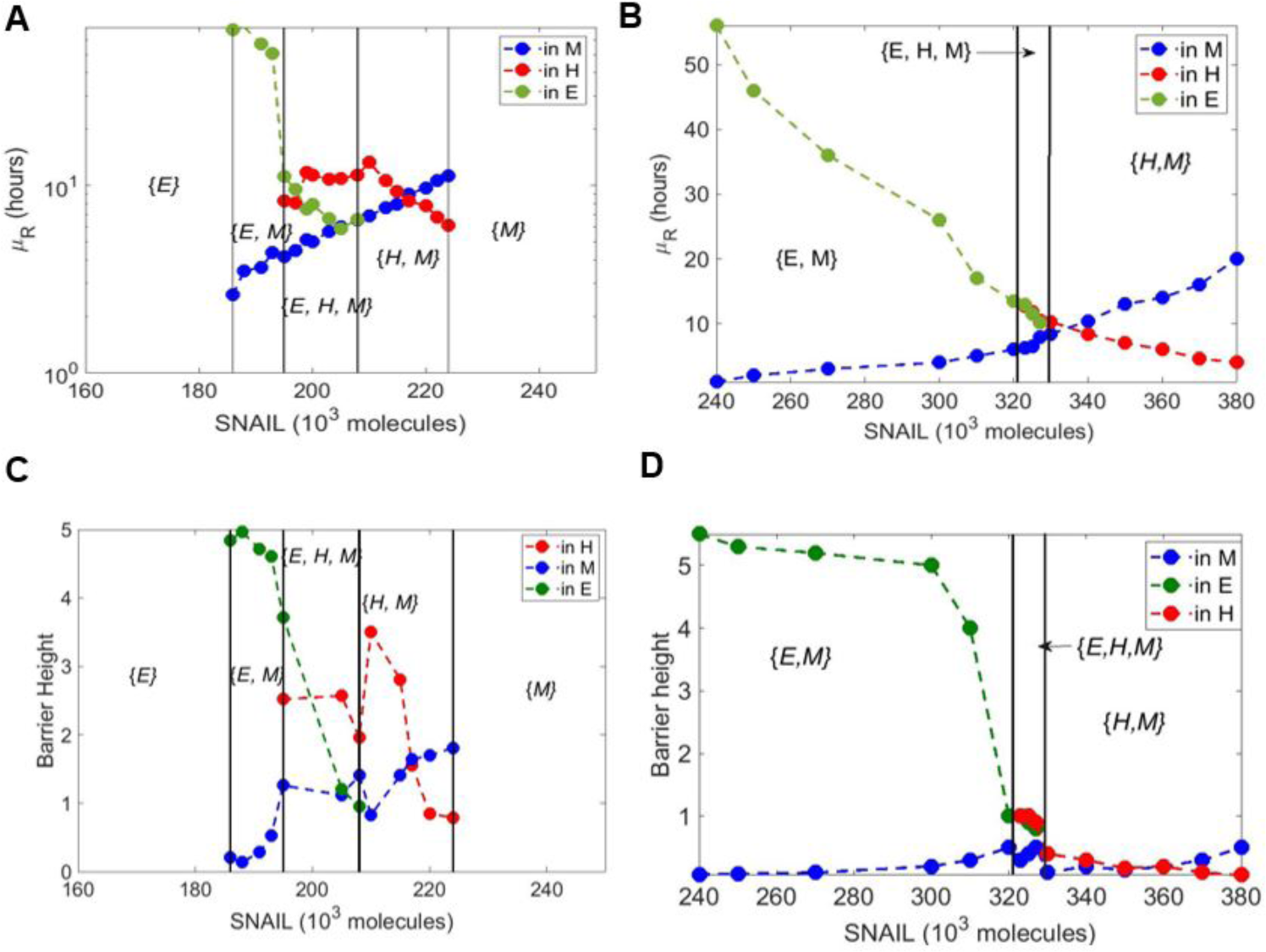
Mean residence time analysis for core EMT circuit vs. coupled EMT-NFATc circuit. A) Variation of mean residence time (µ_R_) with varying SNAIL levels for the core circuit. B) Same as A but for coupled EMT-NFATc circuit. C) Barrier height for core EMT circuit at variable SNAIL levels. D) Same as C but for coupled EMT-NFATc circuit.

### RACIPE analysis of NFATc network reveals its non-canonical behaviour as a PSF

To analyse the underlying design principles of the NFATc-EMT coupled network, we employed a recently developed computational method - Random Circuit Perturbation (RACIPE) (35). RACIPE takes as input the topology of a regulatory network and generates an ensemble of mathematical models corresponding to the network topology, each with a randomly chosen set of kinetic parameters. Then, for each mathematical model, various possible steady states (phenotypes) are identified. Finally, statistical tools are used to identify the robust dynamical properties emerging from the network topology. Here, each mathematical model is a set of five coupled ordinary differential equations (ODEs), where each ODE tracks the temporal dynamics of the five species constituting the regulatory network (SNAIL, ZEB, miR200, E-CAD and NFATc).

Among the 10,000 parameter sets generated via RACIPE, we found cases where the network topology can give rise to the existence of phases with one steady state (mono-stable) or more - two (bi-stable) and three (tri-stable) steady states (Fig 4A). We performed RACIPE on the core EMT network (miR-200/ZEB/SNAIL), its coupling to other PSFs (OVOL, GRHL2, OCT4, NRF2), and the coupled EMT-NFATc network. For a given parameter set, one or more steady states were obtained depending on the initial conditions chosen; each steady state solution was binned as epithelial, hybrid E/M or mesenchymal, based on the z-scores of miR-200 and ZEB for that case. Thus, each parameter was categorized into a given monostable or multistable phase; for instance, a parameter set that enabled both epithelial and hybrid E/M phenotypes for different initial conditions was classified as {E, H} (Fig 4B). Compared to the core network, each of these networks enabled a higher number of parameter sets enabling the co-existence of E and hybrid E/M states ({E, H}) (Fig 4C). Conversely, in most cases, the frequency of {H, M} (co-existence of M and hybrid E/M states) and {E, H, M} (co-existence of E, hybrid E/M and M states) was decreased. However, in the case of NFATc, there was a significant increase in the frequency of {E, H, M} phase (Fig 4C) unlike other PSFs, suggesting that the presence of NFATc may enhance cellular plasticity among epithelial, hybrid E/M and mesenchymal states.

**Fig4:**
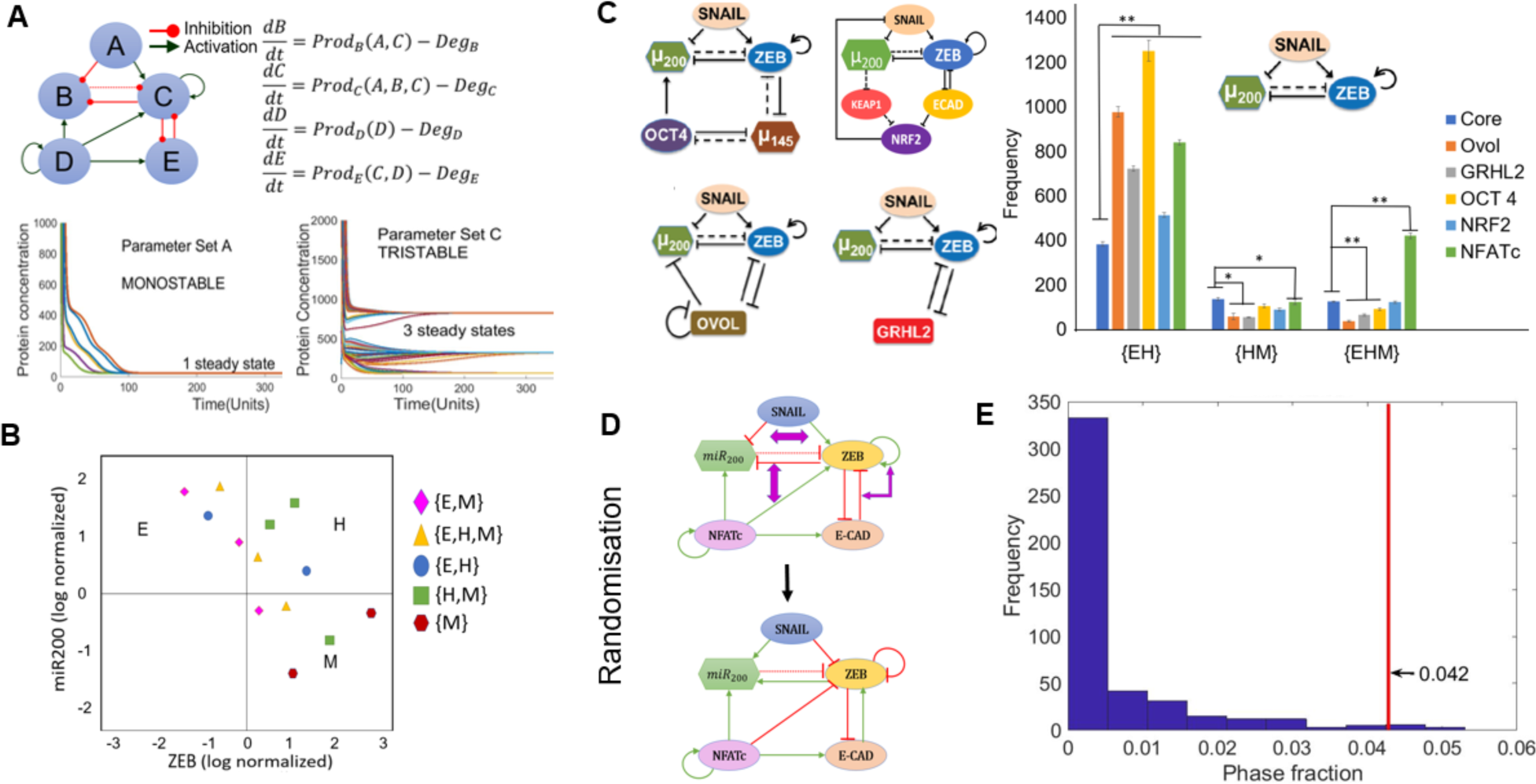
RACIPE analysis of NFATc network. A) Schematic of coupled EMT-NFATc network simulated for various parametric combinations through RACIPE. For different parameters, dynamics of this regulatory network, as modelled by a set of ordinary differential equations representing the interactions in the network, can end up in a mono- or a multi-stable regime. B) Phases are calculated after z-scores of ZEB and miR200 levels calculated from RACIPE are used to classify steady states obtained from each parameter set into Epithelial (E) – (high miR-200, low ZEB)/ Hybrid (H) – (high miR200, high ZEB) / Mesenchymal (M) – (low miR-200, high ZEB) phenotypes. Different symbols denote different phases. C) Frequencies of different multi-stable phases for core EMT network (SNAIL/miR-200/ZEB), its coupling to other PSFs (shown to the left of the graph) and coupled EMT-NFATc coupled network. Error bars denote standard deviation; n=3. D) Depiction of network randomisation methodology. For every node, the degree of incoming and outgoing edges are conserved but not necessarily the number of inhibitory or excitatory edges arriving at or emanating from a node. E) Frequency distribution of the {E,H,M} phase fraction for randomized networks. The red line denotes the phase fraction of {E,H,M} phase in the wild type (WT), i.e. NFATc network. *p < 0.05, **p<0.005 using two tailed Students’ t-test.

To test whether these results for NFATc are specific to the network topology of coupled NTAFc-EMT network, we generated many randomised networks by swapping the edges between nodes in the network, such that the number of incoming and outgoing edges for every node was maintained the same (an example shown in Fig 4D). RACIPE analysis was performed on each of these randomised networks (n=461; see Materials and Methods for details), and the output obtained was separated into different phases i.e. {E}, {H}, {M}, {E, M}, {E, H}, {H, M} and {E, H, M} as mentioned above. We quantified the frequency for multi-stable phases containing the hybrid state i.e. {E, H}, {H, M} and {E, H, M} for all randomized networks, and calculated the frequencies of these phases, i.e. number of parameter sets out of 10,000 that enabled a given phase. The frequency distribution revealed that most of the randomised circuits gave rise to a lower fraction of {E, H, M} phase as compared to that for the wild type NFATc-EMT coupled circuit (the value denoted by the red vertical line) (Fig 4E). However, such stark differences were not observed for the {E, H} (Fig S4A) and {H, M} phases (Fig S4B), suggesting that the NFATc-EMT coupled circuit topology is enriched for enabling co-existence and consequent possible switching among the epithelial, mesenchymal and hybrid E/M phenotypes.

### NFATc confers stability to the hybrid E/M phenotype in a multi-stable phase

We observed that the presence of NFATc in the network increased the frequency of multistable phases containing the hybrid state, particularly the {E, H, M} phase. This led us to investigate the relative stability of the different states in a given multistable phase. To quantify relative stability, every parameter set giving rise to either {E, H}, {H, M} or {E, H, M} phases was simulated using 1000 random initial conditions and each time we tabulated how many initial conditions led to which state – E, H or M. For individual parameter sets, we observed heterogeneity in terms of relative stability of H states in the {E, H} phase, i.e. some parameter sets seemed to have a deeper ‘basin of attraction’ for the epithelial attractor as compared to the hybrid E/M one and *vice versa* (Fig 5A, S5A). Nonetheless, there were similarities in the frequencies of E and the H state obtained across parameter sets obtained from independent RACIPE replicates, as represented by their similar and overlapping kernel density estimates (Fig 5B, S5C-D).

**Fig5:**
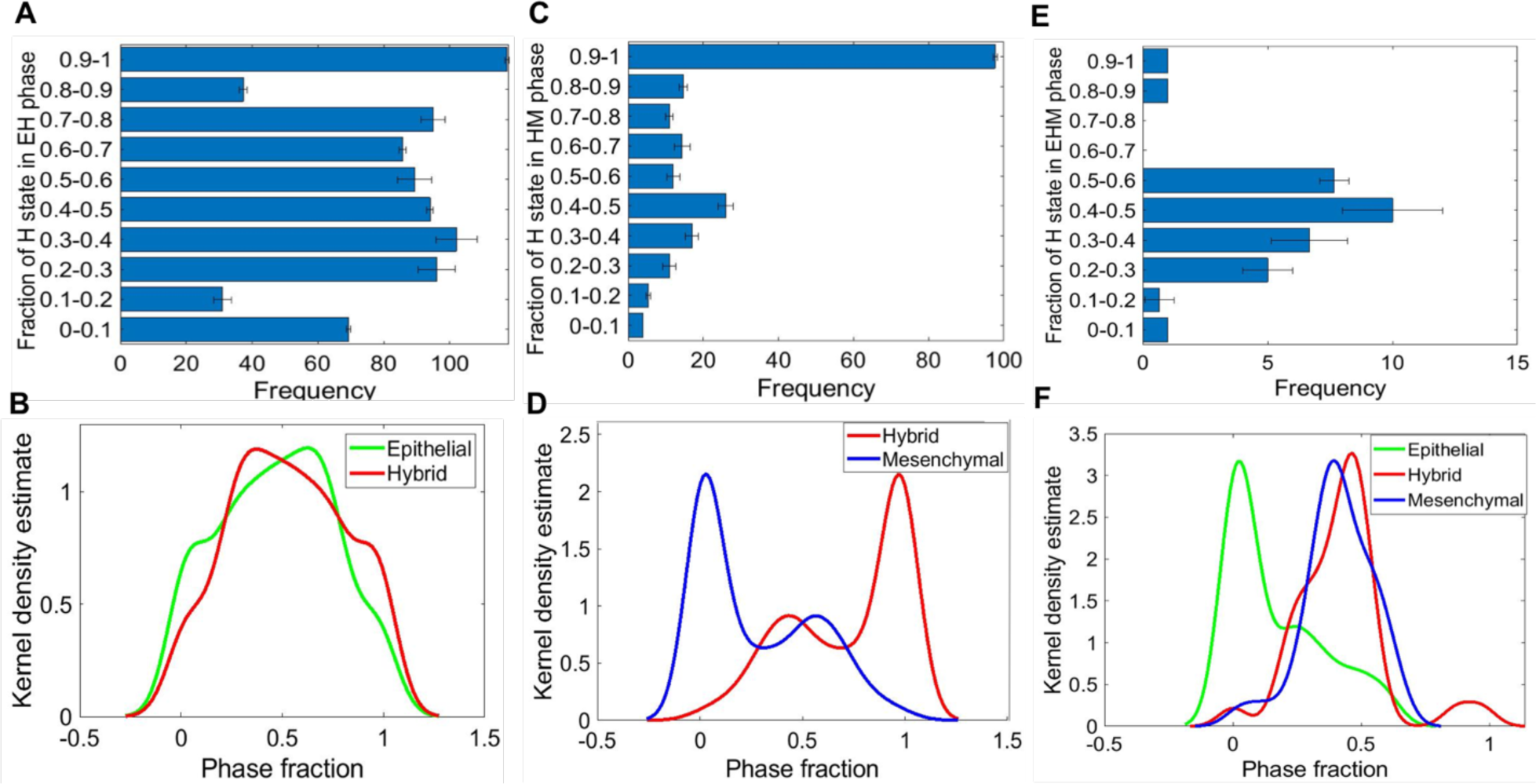
Relative stability analysis. A) Frequency distribution of H state in {E, H} phase. B) Kernel density plot showing the frequency distribution of E and H states in the {E, H} phase. C) Frequency distribution of H state in the {H, M} phase. D) Kernel density plot showing the distribution of M and H states in the {H, M} phase. E) Frequency distribution of H state in the {E, H, M} phase. F) Kernel density plot showing the distribution of E, H and M states in the {E, H, M} phase. For A, C, E; the error bars represent the mean+/- standard deviation for three sets of independently chosen initial conditions for a given parameter set obtained from one RACIPE run.

This analysis for the {H, M} phase revealed that the H state was more stable; i.e. the number of parameter cases for which the relative stability of H state was higher as compared to M state was more than the number of cases when the M state was relatively more stable (Fig 5C,S5B). This trend was maintained for parameter sets obtained across three RACIPE replicates (Fig 5D, S5E, S5F). Similar analysis on the {E, H, M} phase suggested that the E state was relatively less stable than the H and M states (Fig 5E,F, S6A-D). Together, these results indicate that the presence of NFATc can confer high stability to the hybrid E/M state for parametric combinations enabling the co-existence of multiple phenotypes.

### NFATc affects clinical outcome in a tissue specific manner

The hybrid E/M phenotype is often attributed to drive tumour aggressiveness (6,7). This trend is further supported by clinical data where PSFs such as GRHL2 and NRF2 correlate with poor patient survival (17,20). We investigated whether the levels of NFATc can correlate with clinical response, and observed that the association of NFATc with clinical outcomes is context dependent. Higher levels of NFATc correlated with better relapse free survival among breast cancer patients (Fig 6A-C) but with poor relapse free survival among lung cancer patients (Fig 6D). We observed similar context-dependent behaviour of NFATc in terms of overall survival, even within the same tissue (Fig S7A-C). Also, NFATc was positively correlated with better metastasis free survival among breast cancer patients (Fig S7D). Thus, the correlation of NFATc with patient survival is highly likely to be context dependent.

**Fig6:**
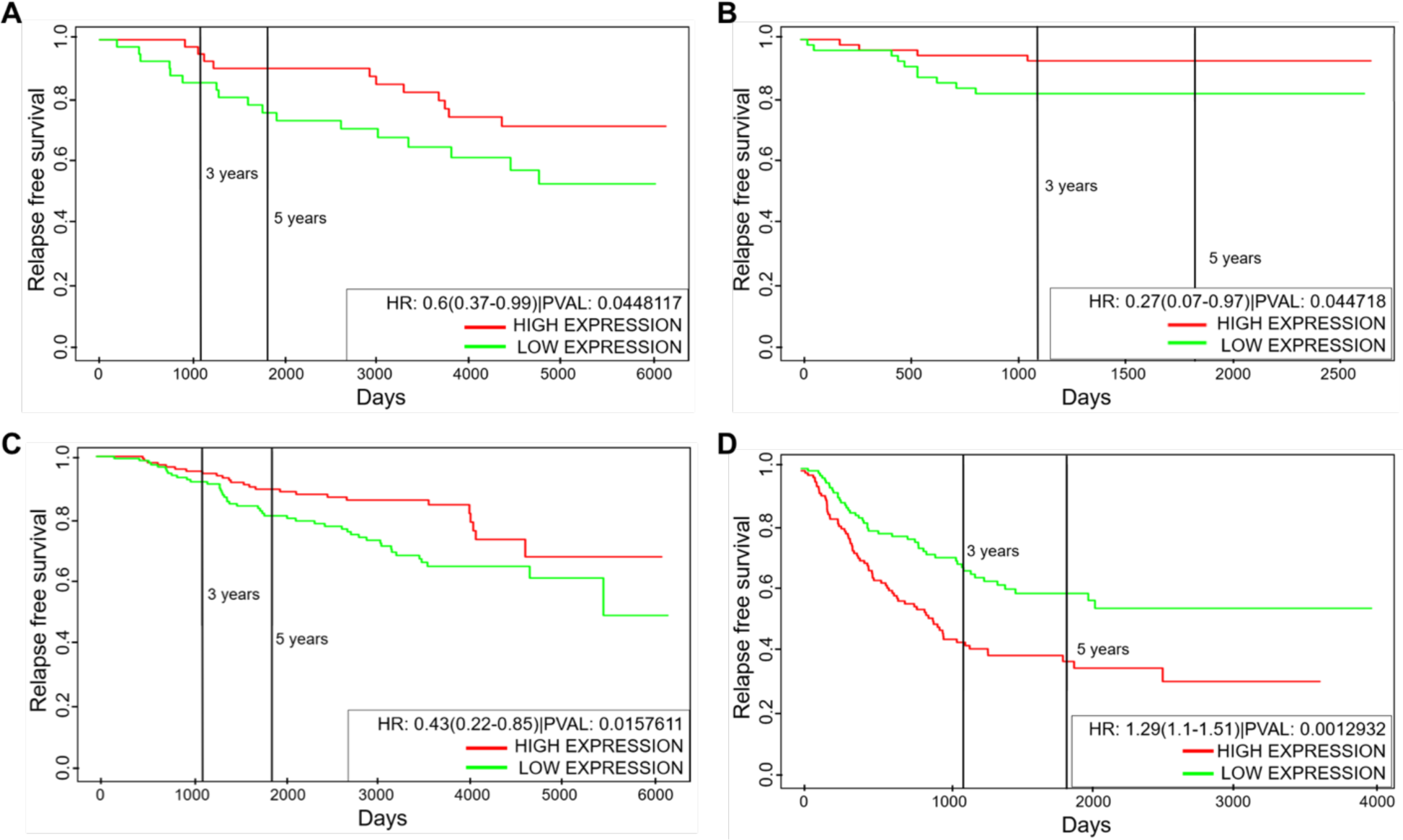
Kaplan-Meier analysis for NFATc1 levels. A-C) Relapse free survival for GSE6532, GSE19615 and GSE17705 (breast cancer). D) Relapse free survival for GSE41271 (lung cancer). Green curve shows the group of patients with low NFATc1; red curve shows the group with high NFATc1 levels. All cohorts are divided at median values of NFATc1; HR denotes hazard ratio; PVAL denotes p-value.

## Discussion

Recent *in vitro, in vivo* and *in silico* investigations have emphasized the existence and significance of hybrid E/M phenotype(s) in various cancer types (36). These hybrid E/M phenotypes can exhibit maximum plasticity (37), possess traits of cancer stem cell-like traits, evade drug resistance, and thus be the ‘fittest’ for metastasis (7). These preclinical experimental observations are supported by clinical analysis of carcinoma samples suggesting that the presence of hybrid E/M cells in a patient at the time of diagnosis associates with poor patient outcomes. Interestingly, even a very small percentage of hybrid E/M cells (score > 2%) was found to be sufficient to confer poor prognosis (38). Thus, identifying mechanisms that can maintain cells in hybrid E/M phenotypes is of crucial importance in our efforts to curb metastatic load.

Here, we developed a computational modelling framework to identify the transcription factor NFATc as a potential phenotypic stability factor (PSF) for hybrid E/M phenotypes. In presence of NFATc, cells undergo a delayed or stalled EMT; thus maintaining cells in hybrid E/M phenotypes; knockdown of NFATc in H1975 NSCLC cells drove the progression towards a complete EMT phenotype, reminiscent of observations made for other PSFs – GRHL2, OVOL2, NUMB and NRF2 (17–22). Similar effects of NFATc knockdown were also seen in MCF10A and DLD1 cells where treatment with VIVIT, a soluble inhibitor of NFATc transcriptional activity, significantly reduced E-cadherin expression and protein level, and increased Slug and Vimentin levels (28), thus driving EMT. NFATc transcriptional activity was shown to be capable of maintaining E-cadherin levels even in the presence of TGFβ induced EMT (28), suggesting that NFATc acted as a “molecular brake” or “guardian” of epithelial traits, preventing a complete EMT (39). Consistently, NFATc1 was identified to be a master regulator of chromatin remodeling to regulate hybrid E/M phenotypes in skin cancer *in vivo*; the proportion of hybrid E/M phenotypes was also shown to be increased by GRHL2, OVOL1/2, and ΔNP63α at the expense of complete EMT cells, thus lending further credence to our results indicating a functional equivalence between NFATc1 and previously identified PSFs such as GRHL2, OVOL1/2, ΔNP63α, and NRF2 (6). In developmental EMT scenario, NFATc1 is implicated in a key role during heart valve development; NFATc1-null embryos exhibit excessive EMT and impaired valve formation. Transcriptional repression of Snail1 and Snail 2 by NFATc1 can inhibit EMT and help maintain vascular E-cadherin levels required for cellular adhesiveness (40,41). These observations across multiple contexts highlight that NFATc may maintain cell-cell contacts in a hybrid E/M phenotype.

Hybrid E/M phenotype(s) are also often associated with higher stem-like behaviour and enhanced metastasis across cancer types *in vitro* and *in vivo* (24). Consistently, NFAT transcriptional activity contributes to metastasis in colon cancer; inhibition of NFATc1 reduced metastatic growth in an immunocompetent mouse model. Further, genes upregulated by NFATc1 significantly correlated with worse clinical outcomes for Stage II and III colorectal cancer patients (42). Similarly, NFATc2 was overexpressed in lung adenocarcinoma tumour-initiating cells; it supported tumorigenesis *in vivo* and its knockdown *in vitro* reduced 60-70% tumor-spheres and restricted the renewability of tumor-spheres (31). NFAT/calcineurin signaling pathway is also activated in breast cancer and aggravates tumorigenic and metastatic potential of mammary tumour cells *in vitro* and *in vivo* (43,44). Furthermore, NFATc1 levels were found to be significantly upregulated in spheroid-forming cells in pancreatic cancer, where NFATc1 promotes SOX2 transcriptionally (27). One of the targets of NFAT/SOX2 signaling pathway is ALDH1A1(31) – a *bona fide* marker for hybrid E/M cells behaving as cancer stem cells in breast cancer (45). Therefore, these observations underscore the connection between NFAT signaling, stemness, and metastatic aggressiveness.

Our results show that while NFATc increased the parametric range of SNAIL levels enabling a hybrid E/M phenotype, it did not increase the mean residence time of hybrid E/M cells, suggesting that the role of NFATc may be non-canonical in terms of behaving as a PSF. This non-canonical behaviour is further elucidated by RACIPE analysis, where, unlike other PSFs such as GRHL2, NFATc increased the frequency of parametric combinations containing co-existing epithelial, hybrid E/M and M phenotypes, and possible interconversions among them. Thus, NFATc may be thought of as a driver of phenotypic plasticity, and targeting NFAT signaling may curb cancer cell adaptation (46) - a distinctive property of metastasis-initiating cells (47).

## Acknowledgements

This work was supported by Ramanujan Fellowship awarded by SERB, DST, Government of India to MKJ (SB/S2/RJN-049/2018).

## Author contributions

MKJ and SCT designed research; MKJ, SCT, AG and SH supervised research; ARS, DN, and KB performed research; all authors contributed to analysing data and writing and editing of the manuscript.

## Materials and Methods

### Mathematical Modelling

As per the schematic shown in Fig 1A, the dynamics of all five molecular species (miR-200, Snail, Zeb, E-cadherin and NFATc) was described by a system of coupled ordinary differential equations (ODEs). The level of a protein, mRNA or micro-RNA (X) is described via a chemical rate equation that assumes the generic form:

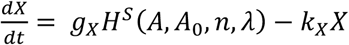

Where the first term of the equation signifies the basal rate of production (*g*_*X*_); the terms multiplied to *g*_*X*_ represent the transcriptional/translational/post-translational regulations due to interactions among the species in the system, as defined by the Hills function (*H*^*S*^(*A, A*_0_, *n, λ*)). The term *k*_*X*_*X* accounts for the rate of degradation of the species (X) based on first order kinetics. The complete set of equations and parameters are presented in the SI.

### Cell culture and siRNA treatments

H1975 cells were cultured in RPMI 1640 medium containing 10% fatal bovine serum and 1% penicillin/streptomycin cocktail (Thermo Fisher Scientific). Cells were transfected at a final concentration of 50nM siRNA using Lipofectamine RNAiMAX (Thermo Fisher Scientific) according to the manufacturer’s instructions using following siRNAs: siControl (Thermo Fisher Scientific), siNFATC1 #1 (Invitrogen), siNFATC1 #2 (Invitrogen). Regular mycoplasma testing was also carried out to exclude any possible cell culture contamination.

### RT-PCR

Total RNA was isolated following manufacturer’s instructions using RNAeasy kit (Qiagen). cDNA was prepared using iScript gDNA clear cDNA synthesis kit (Bio-Rad). A TaqMan PCR assay was performed with a 7500 Fast Real-Time PCR System using TaqMan PCR master mix, commercially available primers, and FAM^™^-labelled probes for CDH1, VIM, ZEB1, NFATC1, SNAIL, and VIC^™^-labelled probes for 18 S, as per manufacturer’s instructions (Life Technologies). Each sample was run in biological and technical triplicates. Ct values for each gene was calculated and normalized to Ct values for 18 S (ΔCt). The ΔΔCt values were then calculated by normalization to the ΔCt value for control.

### Western blotting analysis

H1975 cells were lysed in RIPA lysis assay buffer (Pierce) supplemented with enzyme inhibitor cocktail (Roche). The samples were separated on a 4–20% SDS-polyacrylamide gel (Biorad). After transfer to PVDF membrane, incubation was carried out with primary antibodies anti-CDH1 (1:1000; Cell Signaling Technology), anti-vimentin (1:1000; Cell Signaling Technology), anti-Zeb1 (1:1000; Cell Signaling Technology), anti-SNAIL (1:1000; Cell Signaling Technology), and anti-beta actin (1:10 000; Abcam) and subsequent secondary antibodies. Membranes were exposed using the ECL method (GE Healthcare) as per manufacturer’s instructions.

### Immunofluorescence

H1975 Cells were fixed in 3.4% paraformaldehyde, permeabilized with 0.2% Triton X-100, and then stained with primary antibodies against CDH1 (1:100; Abcam) and vimentin (1:100; Cell Signaling Technology). Alexa conjugated secondary antibodies (Life technologies) were used to detect the expression of respective proteins. DAPI was used to counterstain the nuclei.

### Wound-Healing assay

Scratch wound-healing assay was performed to determine cell migration using confluent cultures (80-90% confluence). Briefly, H1975 cells (1 × 105 cells/ml) were seeded in 6-well tissue culture plate. After cells attain expected confluency, they were starved for 24 hours using 0.2% serum in growth media. Next day, a sterile p200 pipet tip was used to create a wound on the confluent monolayer and media was replenished. Images were acquired at 0 and 16 hours; the experiments were repeated 3 times. Images of the scratch wounds were taken and measured by Image-J software to calculate the mean and standard deviation. Each group was compared with the control group. Cell migration was expressed as the migration rate: (original scratch width - new scratch width)/original scratch width × 100%.

### Trans-well migration assay

H1975 cells were grown in 6-well plates and treated with siNFATC1 for 24 hours. After 48hrs of NFATC1 knockdown, cell monolayers were harvested, and 2 × 104 viable cells/200 µl cell concentration was prepared in serum free medium. The cell suspension was transferred on top of a 0.8 µm pore diameter Transwell insert (Millipore) and placed on a 24 well cell culture plate. A 10% fetal bovine serum solution was added as chemo-attractant at the bottom of the insert and plate was incubated at 37°C for 18 hours. Non-migrated cells were removed by gently swabbing inside each insert. Cells were fixed and stained with a 0.5% crystal violet solution for 10 minutes. The inserts were thoroughly washed with and air dried completely before visualizing under a microscope. Cell numbers were counted at ×200 magnification. The experiment was repeated 3 times and statistically analysed with 5 fields of view, and the mean values were taken as the migratory cell number.

### Mean residence time analysis

The mean residence time was calculated as follows: As the degradation rate of ZEB mRNA is much greater than that of ZEB protein and miR-200 and also the production rate of E-cad and NFATc is much more larger than that of ZEB protein and miR-200, we assumed that ZEB mRNA, E-cad, and NFATc reach to the equilibrium much faster relatively, that is 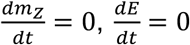, and 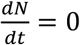. This assumption reduces the equations given in SI: to two coupled ODEs of ZEB and miR-200. Then we simulated the dynamical system in presence of external noise and obtained the time evolution of ZEB protein and miR-200 using Euler-Maruyama simulation. From the time evolution of ZEB and miR-200 the dynamical states of the system were coarse grained as an itinerary of basins visited. Then the MRT was calculated by multiplying the total number of successive states with Δt. Detailed methods are outlined in the publication by Biswas *et al.*(23).

### RACIPE

Random circuit perturbation (RACIPE) (35) algorithm was run on the coupled EMT-NFATc network and its randomized counterparts. Continuous steady state levels were obtained as output for the five variables, for ensembles of mathematical models; each model has a randomly chosen parameter set corresponding to intrinsic production/degradation of all species as well as those representing the regulatory links. The algorithm was used to generate 10,000 mathematical models, each with a different set of parameters. 100 initial conditions were chosen for each model, and all steady state solutions obtained were compiled together. With this consolidated data, the z-scores of steady state levels of all the biomolecules in the individual networks were calculated. Based on the z-score of ZEB and miR-200, the phenotype for a given steady state solution is decided, i.e. if (z_zeb_>0) and (z_mir-200_<0), it is counted as mesenchymal state, (z_zeb_>0) and (z_mir-200_>0) is counted as a hybrid state, and (z_zeb_<0) and (z_mir-200_>0) is counted as an epithelial state. Similarly, states are determined for all solutions and based on the state of each steady state for a given set of parameters, the phases are determined.

### Network Randomisation

The following rules were employed to generate an ensemble of randomized networks. For each node, in each instance of a randomization of the wild type network (FIG 1A), the number of incoming and outgoing edges were kept constant. The number of activation edges and the number of inhibitory edges were also kept fixed at 6 and 5 respectively (The same number as that in the wild type network (FIG 1A)). Furthermore, the source node and the target node for each of the edges were kept fixed but the identity of the edge in terms of it being an activation or inhibition link was allowed to change. Hence 461 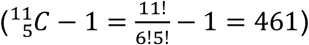 such randomized networks were constructed excluding the wild type case.

### Relative stability analysis

The relative stability analysis was performed in MATLAB. RACIPE generated parameter sets that give rise to multi-stable phases were first determined. These parameter sets were simulated in MATLAB using 1000 random initial conditions and the steady state each time was determined. Then, the total number of times each of the possible steady states was reached was calculated.

### Kaplan-Meier analysis

ProgGene (48) was used for conducting Kaplan-Meier analysis for respective datasets. The number of samples in NFATc1-high vs. NFATc1-low categories are given below:

GSE6532 (breast cancer): n(High) = 44, n(Low)= 43
GSE19615 (breast cancer) : n(High) = 58, n(Low) = 57
GSE17705 (lung cancer): n(High)= 149, n (Low)=149
GSE41271 (lung cancer): n(High) = 138, n(Low) = 137
GSE30219 (lung cancer): n (High) = 141 n (Low) = 141
GSE19536 (breast cancer): n(High)- 56, n(Low) = 44
GSE14814 (lung cancer): n (High) = 44, n (Low) = 44:

## References

1. Gupta GP, Massagué J. Cancer metastasis: building a framework. Cell (2006) 127:679–695.

2. Jolly MK, Tripathi SC, Somarelli JA, Hanash SM, Levine H. Epithelial-mesenchymal plasticity : How have quantitative mathematical models helped improve our understanding ? Mol Oncol (2017) 11:739–754. doi: 10.1002/1878-0261.12084

3. Jolly MK, Kulkarni P, Weninger K, Orban J, Levine H. Phenotypic plasticity, bet-hedging, and androgen independence in prostate cancer: Role of non-genetic heterogeneity. Front Oncol (2018) 8:50. doi: 10.3389/fonc.2018.00050

4. Gupta PB, Fillmore CM, Jiang G, Shapira SD, Tao K, Kuperwasser C, Lander ES. Stochastic state transitions give rise to phenotypic equilibrium in populations of cancer cells. Cell (2011) 146:633–644. doi: 10.1016/j.cell.2011.07.026

5. Nieto MA, Huang RY, Jackson RA, Thiery JP. EMT: 2016. Cell (2016) 166:21–45.

6. Pastushenko I, Blanpain C. EMT Transition States during Tumor Progression and Metastasis. Trends Cell Biol (2019) 29:212–226. doi: 10.1016/j.tcb.2018.12.001

7. Jolly MK, Mani SA, Levine H. Hybrid epithelial/mesenchymal phenotype(s): The ‘fittest’ for metastasis? Biochim Biophys Acta - Rev Cancer (2018) 1870:151–157. doi: 10.1016/j.bbcan.2018.07.001

8. Campbell K, Rossi F, Adams J, Pitsidianaki I, Barriga FM, Garcia-Gerique L, Batlle E, Casanova J, Casali A. Collective cell migration and metastases induced by an epithelial-to-mesenchymal transition in Drosophila intestinal tumors. Nat Commun (2019) 10:2311. doi: 10.1038/s41467-019-10269-y

9. Liao T-T, Yang M-H. Hybrid Epithelial/Mesenchymal State in Cancer Metastasis: Clinical Significance and Regulatory Mechanisms. Cells (2020) 9:623.

10. Bocci F, Jolly MK, Onuchic JN. A biophysical model uncovers the size distribution of migrating cell clusters across cancer types. Cancer Res (2019) 79:5527–5535. doi: 10.1158/0008-5472.CAN-19-1726

11. Giuliano M, Shaikh A, Lo HC, Arpino G, De Placido S, Zhang XH, Cristofanilli M, Schiff R, Trivedi M V. Perspective on Circulating Tumor Cell Clusters: Why It Takes a Village to Metastasize. Cancer Res (2018) 78:845–852. doi: 10.1158/0008-5472.CAN-17-2748

12. Gonzalez DM, Medici D. Signaling mechanisms of the epithelial-mesenchymal transition. (2014) 7:1–17. doi: 10.1126/scisignal.2005189

13. Burk U, Schubert J, Wellner U, Schmalhofer O, Vincan E, Spaderna S, Brabletz T. A reciprocal repression between ZEB1 and members of the miR-200 family promotes EMT and invasion in cancer cells. EMBO Rep (2008) 9:582–589. doi: 10.1038/embor.2008.74

14. Lu M, Jolly MK, Levine H, Onuchic JN, Ben-Jacob E. MicroRNA-based regulation of epithelial-hybrid-mesenchymal fate determination. Proc Natl Acad Sci U S A (2013) 110:18174–9. doi: 10.1073/pnas.1318192110

15. Siemens H, Jackstadt R, Hünten S, Kaller M, Menssen A, Götz U, Hermeking H. miR-34 and SNAIL form a double-negative feedback loop to regulate epithelial-mesenchymal transitions. Cell Cycle (2011) 10:4256–4271. doi: 10.4161/cc.10.24.18552

16. Park S-MM, Gaur AB, Lengyel E, Peter ME. The miR-200 family determines the epithelial phenotype of cancer cells by targeting the E-cadherin repressors ZEB1 and ZEB2. Genes Dev (2008) 22:894–907. doi: 10.1101/gad.1640608

17. Bocci F, Tripathi SC, Vilchez MSA, George JT, Casabar J, Wong P, Hanash S, Levine H, Onuchic J, Jolly M. NRF2 activates a partial Epithelial-Mesenchymal Transition and is maximally present in a hybrid Epithelial/Mesenchymal phenotype. Integr Biol (2019) 11:251–263. doi: 10.1101/390237

18. Watanabe K, Villarreal-Ponce A, Sun P, Salmans ML, Fallahi M, Andersen B, Dai X. Mammary morphogenesis and regeneration require the inhibition of EMT at terminal end buds by Ovol2 transcriptional repressor. Dev Cell (2014) 29:59–74. doi: 10.1016/j.devcel.2014.03.006

19. Jia D, Jolly MK, Boareto M, Parsana P, Mooney SM, Pienta KJ, Levine H, Ben-Jacob E. OVOL guides the epithelial-hybrid-mesenchymal transition. Oncotarget (2015) 6:15436–48. doi: 10.18632/oncotarget.3623

20. Jolly MK, Tripathi SC, Jia D, Mooney SM, Celiktas M, Hanash SM, Mani SA, Pienta KJ, Ben-Jacob E, Levine H. Stability of the hybrid epithelial/mesenchymal phentoype. Oncotarget (2016) 7:27067–27084.

21. Bocci F, Jolly MK, Tripathi SC, Aguilar M, Hanash SM, Levine H, Onuchic JN. Numb prevents a complete epithelial-mesenchymal transition by modulating Notch signaling. J R Soc Interface (2017) 14: doi: 10.1098/rsif.2017.0512

22. Hong T, Watanabe K, Ta CH, Villarreal-Ponce A, Nie Q, Dai X. An Ovol2-Zeb1 Mutual Inhibitory Circuit Governs Bidirectional and Multi-step Transition between Epithelial and Mesenchymal States. PLOS Comput Biol (2015) 11:e1004569. doi: 10.1371/journal.pcbi.1004569

23. Biswas K, Jolly M, Ghosh A. Stability and mean residence times for hybrid epithelial/mesenchymal phenotype. Phys Biol (2019) 16:025003. doi: 10.1088/1478-3975/aaf7b7

24. Jolly MK, Somarelli JA, Sheth M, Biddle A, Tripathi SC, Armstrong AJ, Hanash SM, Bapat SA, Rangarajan A, Levine H. Hybrid epithelial/mesenchymal phenotypes promote metastasis and therapy resistance across carcinomas. Pharmacol Ther (2019) 194:161–184. doi: 10.1016/j.pharmthera.2018.09.007

25. Mognol GP, Carneiro FRG, Robbs BK, Faget D V., Viola JPB. Cell cycle and apoptosis regulation by NFAT transcription factors: New roles for an old player. Cell Death Dis (2016) 7: doi: 10.1038/cddis.2016.97

26. Medyouf H, Ghysdael J. The calcineurin/NFAT signaling pathway: A novel therapeutic target in leukemia and solid tumors. Cell Cycle (2008) 7:297–303. doi: 10.4161/cc.7.3.5357

27. Singh SK, Chen N, Hessmann E, Siveke J, Lahmann M, Singh G, Voelker N, Vogt S, Esposito I, Schmidt A, et al. Antithetical NFAT c1–Sox2 and p53–miR200 signaling networks govern pancreatic cancer cell plasticity. EMBO J (2015) 34:517–530. doi: 10.15252/embj.201489574

28. Gould R, Bassen DM, Chakrabarti A, Varner JD, Butcher J. Population Heterogeneity in the Epithelial to Mesenchymal Transition Is Controlled by NFAT and Phosphorylated Sp1. PLoS Comput Biol (2016) 12:e005251. doi: 10.1371/journal.pcbi.1005251

29. Schmalhofer O, Brabletz S, Brabletz T. E-cadherin, beta-catenin, and ZEB1 in malignant progression of cancer. Cancer Metastasis Rev (2009) 28:151–66. doi: 10.1007/s10555-008-9179-y

30. Mooney SM, Jolly MK, Levine H, Kulkarni P. Phenotypic plasticity in prostate cancer: role of intrinsically disordered proteins. Asian J Androl (2016) 18:704–10. doi: 10.4103/1008-682X.183570

31. Xiao ZJ, Liu J, Wang SQ, Zhu Y, Gao XY, Tin VPC, Qin J, Wang JW, Wong MP. NFATc2 enhances tumor-initiating phenotypes through the NFATc2/SOX2/ALDH axis in lung adenocarcinoma. Elife (2017) 6: doi: 10.7554/eLife.26733

32. Wang G, Guo X, Hong W, Liu Q, Wei T, Lu C, Gao L, Ye D, Zhou Y. Critical regulation of miR-200 / ZEB2 pathway in Oct4 / Sox2-induced mesenchymal-to-epithelial transition and induced pluripotent stem cell generation. Proc Natl Acad Sci U S A (2013) 110:2858–63. doi: 10.1073/pnas.1212769110

33. Somarelli JA, Shelter S, Jolly MK, Wang X, Bartholf Dewitt S, Hish AJ, Gilja S, Eward WC, Ware KE, Levine H, et al. Mesenchymal-epithelial transition in sarcomas is controlled by the combinatorial expression of miR-200s and GRHL2. Mol Cell Biol (2016) 36:2503–13. doi: 10.1128/MCB.00373-16

34. Jolly MK, Boareto M, Debeb BG, Aceto N, Farach-Carson MC, Woodward WA, Levine H. Inflammatory Breast Cancer: a model for investigating cluster-based dissemination. NPJ Breast Cancer (2017) 3:21. doi: https://doi.org/10.1101/119479

35. Huang B, Lu M, Jia D, Ben-Jacob E, Levine H, Onuchic JN. Interrogating the topological robustness of gene regulatory circuits by randomization. PLoS Comput Biol (2017) 13:e1005456. doi: 10.137/journal.pcbi.1005456

36. Jolly MK, Celia-Terrassa T. Dynamics of Phenotypic Heterogeneity Associated with EMT and Stemness during Cancer Progression. J Clin Med (2019) 8:1542. doi: 10.3390/jcm8101542

37. Tripathi S, Chakraborty P, Levine H, Jolly MK. A mechanism for epithelial-mesenchymal heterogeneity in a population of cancer cells. PLoS Comput Biol (2020) 16:e1007619. doi: 10.1101/592691

38. Godin L, Balsat C, Eycke Y Van, Allard J, Royer C, Remmelink M, Pastushenko I, Haene ND, Blanpain C, Salmon I, et al. A Novel Approach for Quantifying Cancer Cells Showing Hybrid Epithelial / Mesenchymal States in Large Series of Tissue Samples : Towards a New Prognostic Marker. Cancers (Basel) (2020) 12:906. doi: 10.3390/cancers12040906

39. Li S, Yang J. Ovol Proteins: Guardians against EMT during Epithelial Differentiation. Dev Cell (2014) 29:1–2. doi: 10.1016/j.devcel.2014.04.002

40. Wu B, Baldwin HS, Zhou B. Nfatc1 directs the endocardial progenitor cells to make heart valve primordium. Trends Cardiovasc Med (2013) 23:294–300. doi: 10.1016/j.tcm.2013.04.003

41. Wu B, Wang Y, Lui W, Langworthy M, Tompkins KL, Hatzopoulos AK, Baldwin HS, Zhou B. Nfatc1 coordinates valve endocardial cell lineage development required for heart valve formation. Circ Res (2011) 109:183–92. doi: 10.1161/CIRCRESAHA.111.245035

42. Tripathi MK, Deane NG, Zhu J, An H, Mima S, Wang X, Padmanabhan S, Shi Z, Prodduturi N, Ciombor KK, et al. Nuclear factor of activated T-cell activity is associated with metastatic capacity in colon cancer. Cancer Res (2014) 74:6947–6957. doi: 10.1158/0008-5472.CAN-14-1592

43. Quang CT, Lebouche S, Passaro D, Furhmann L, Nourieh M, Vincent-Salomon A, Ghysdael J. The calcineurin/NFAT pathway is activated in diagnostic breast cancer cases and is essential to survival and metastasis of mammary cancer cells. Cell Death Dis (2015) 6:e1658. doi: 10.1038/cddis.2015.14

44. Yiu GK, Toker A. NFAT induces breast cancer cell invasion by promoting the induction of cyclooxygenase-2. J Biol Chem (2006) 281:12210–12217. doi: 10.1074/jbc.M600184200

45. Colacino JA, Azizi E, Brooks MD, Harouaka R, Fouladdel S, McDermott SP, Lee M, Hill D, Madde J, Boerner J, et al. Heterogeneity of human breast stem and progenitor cells as revelaed by transcriptional profiling. Stem Cell Reports (2018) 10:1596–1609. doi: 10.1016/j.stemcr.2016.05.008

46. Qin J-J, Nag S, Wang W, Zhou J, Zhang W-D, Wang H, Zhang R. NFAT as cancer target: Mission possible? Biochim Biophys Acta (2014) 1846:297–311. doi: 10.1016/j.bbcan.2014.07.009

47. Celià-Terrassa T, Kang Y. Distinctive properties of metastasis-initiating cells. Genes Dev (2016) 30:892–908. doi: 10.1101/gad.277681.116

48. Goswami CP, Nakshatri H. PROGgeneV2 : enhancements on the existing database. BMC Cancer (2014) 14:

