## Supplementary Information for "NFATc acts as a non-canonical phenotypic stability factor for a hybrid epithelial/mesenchymal phenotype"

### Mathematical modelling

The dynamics of the NFATc-EMT coupled circuit describes the dynamics of the molecular species of the EMT regulatory circuit (miR-200, Snail, Zeb), E-cadherin and NFATc as shown in Fig 1A. This set-up extends the mathematical model of EMT circuit previously developed (1). The given set of coupled ordinary differential equations (ODEs) represent the dynamics of the species of the circuit (miR-200: $\mu_{200}$, Snail: S, Zeb: Z, E-cadherin: E, NFATc: N):

$$\frac{d\mu_{200}}{dt} = g_{\mu_{200}} H^s(Z,\lambda_{Z,\mu_{200}}) H^s(S,\lambda_{S,\mu_{200}}) H^s(N,\lambda_{N,\mu_{200}}) - m_Z Y_\mu(\mu_{200}) - k_{\mu_{200}}\mu_{200}$$

$$\frac{dm_Z}{dt} = g_{m_Z} H^s(Z,\lambda_{Z,m_Z}) H^s(S,\lambda_{S,m_Z}) H^s(N,\lambda_{N,m_Z}) H^s(E,\lambda_{E,m_Z}) - m_Z Y_Z(\mu_{200}) - k_{m_Z} m_Z$$

$$\frac{dZ}{dt} = g_Z m_Z L(\mu_{200}) - k_Z Z$$

$$\frac{dE}{dt} = g_E H^s(Z,\lambda_{Z,E}) H^s(N,\lambda_{N,E}) - k_E E$$

$$\frac{dN}{dt} = g_N H^s(N,\lambda_{N,N}) - k_N N$$

where $g_X$ is the corresponding innate production rate and $k_X$ is the innate degradation rate.

$m_z L(\mu_{200})$ is the net translation rate, $m_z Y_m(\mu_{200})$ is the total ZEB mRNA active degradation rate and $m_z Y_\mu(\mu_{200})$ is the total miR active degradation rate. $H^S$ is the shifted Hill function, defined as

$H^S(B,\lambda) = H^-(B)+\lambda H^+(B)$ ,

$H^-(B) = 1/[1+(B/B_0)^{nB}]$ ,

$H^+(B) = 1-H^-(B)$,

λ is the fold change from the basal synthesis rate due to protein B. λ >1 for activators, while λ<1 for inhibitors.

### Parameter estimation

The parameters were adopted from previously published literature for the molecular species of the EMT regulatory circuit (miR-200, Snail, Zeb), E-cadherin and NFATc interactions.

| Parameter | Value | Reference |
|---|---|---|
| $g\mu200$ (Molecules/Hour) | 2.1K | (1) |
| gmZ (Molecules/Hour) | 11 | (1) |
| $Z^0\mu_{200}$ (Molecules) | 220K | (1) |
| $Z^0 m_Z$ (Molecules) | 25K | (1) |
| $n_{Z,\mu_{200}}$ | 3 | (1) |
| $n_{Z,m_Z}$ | 2 | (1) |
| $n_{\mu_{200}}$ | 6 | (1) |
| $n_{S,\mu_{200}}$ | 2 | (1) |
| $n_{S,m_Z}$ | 2 | (1) |
| $\lambda_{Z,\mu_{200}}$ | 0.1 | (1) |
| $\lambda_{Z,m_Z}$ | 7.5 | (1) |
| $\lambda_{S,\mu_{200}}$ | 0.1 | (1) |
| $\lambda_{S,m_Z}$ | 10 | (1) |
| $k_{\mu_{200}}$(Hour$^{-1}$) | 0.05 | (1) |

| | | |
|---|---|---|
| $k_{m_Z}$(Hour$^{-1}$) | 0.5 | (1) |
| $k_Z$(Hour$^{-1}$) | 0.1 | (1) |
| $g_Z$(Hour$^{-1}$) | 0.1K | (1) |
| $S^0_{\mu_{200}}$(Molecules) | 180K | (1) |
| $S^0_{m_Z}$(Molecules) | 180K | (1) |
| S(Molecules) | 200K | (1) |
| $\mu^0_{200}$(Molecules) | 10K | (1) |
| $g_E$(Molecules/Hour) | 5000 | (2) |
| $k_E$(Hour$^{-1}$) | 0.1 | (2) |
| $\lambda_{E,m_Z}$ | 0.8 | (2) |
| $n_{E,m_Z}$ | 2 | (2) |
| $E^0_{m_Z}$(Molecules) | 80000 | (2) |
| $\lambda_{Z,E}$ | 0.1 | (2) |
| $n_{Z,E}$ | 2 | (2) |
| $Z^0_E$ | 100000 | (2) |
| $\lambda_{N,E}$ | 8 | (3) |
| $n_{N,E}$ | 5 | (4) |
| $N^0_E$ | 100000 | Estimated |
| $g_N$(Molecules/Hour) | 80000 | (4) |
| $k_N$(Hour$^{-1}$) | 0.1 | (4) |
| $\lambda_{N,Z}$ | 2 | (4) |
| $n_{N,Z}$ | 2 | (4) |
| $N^0_Z$ | 800000 | Estimated |
| $\lambda_{N,\mu_{200}}$ | 4 | Estimated |
| $n_{N,\mu_{200}}$ | 4 | Estimated |
| $N^0_{\mu 200}$ (Molecules) | 500000 | Estimated |
| $\lambda_{N,N}$ | 7 | (5) |
| $n_{N,N}$ | 3 | (5) |
| $N^0_N$ | 100000 | Estimated |

K denotes $10^3$ molecules

**Kaplan Meier analysis**

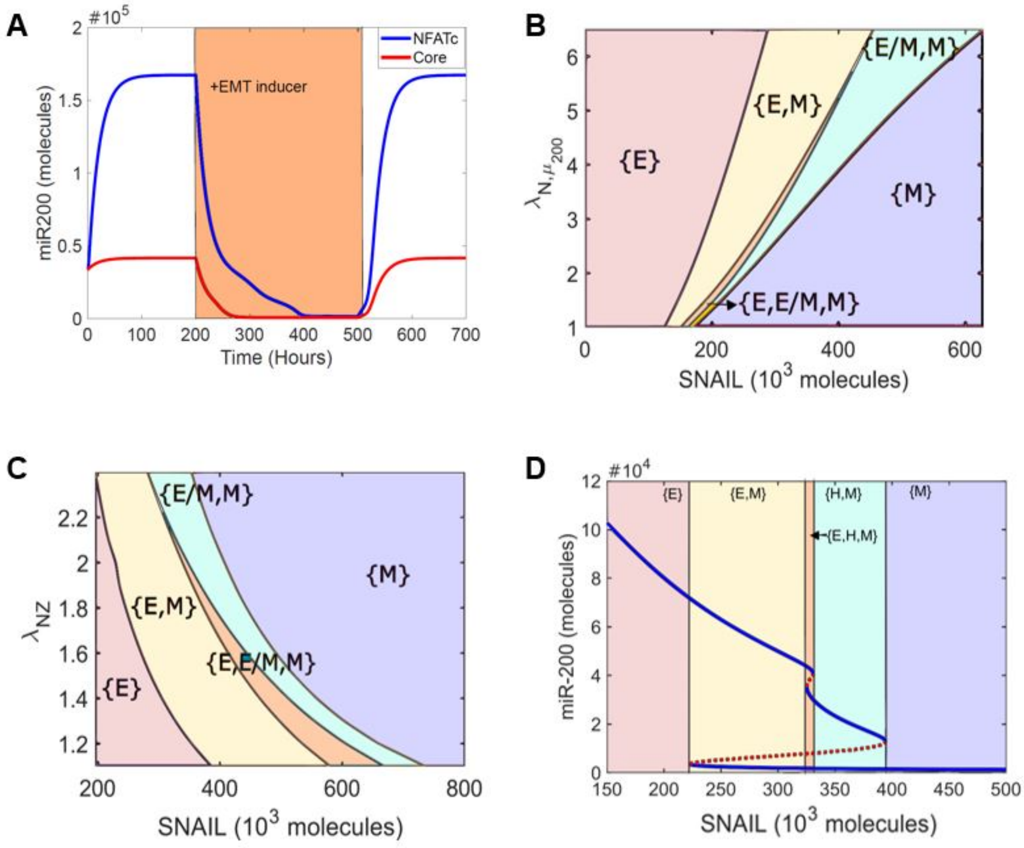


***FigS1: NFATc inhibits a complete EMT*** *A) Dynamics of miR200 levels in a cell starting in an epithelial phenotype, when exposed to a high level of S=330000 molecules (orange-shaded region). (B) Phase diagram of NFATc network when driven by SNAIL and varying strength of interaction between NFATc and miR200. (C) Phase diagram of NFATc network when driven by SNAIL and varying strength of interaction between NFATc and ZEB. (D) Bifurcation diagram of miR200 levels as driven by SNAIL signal.*

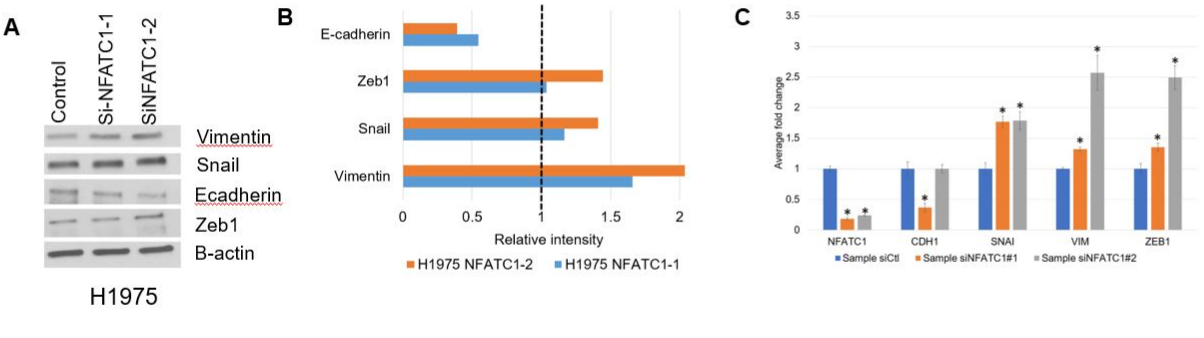


***FigS2: Knockdown of NFATc drives a complete EMT.*** *A) Quantification of western blot analysis shows decreased levels of E-CAD and increased levels of VIM, SNAIL and ZEB with knockdown of NFATc1. (B) Quantitative RT-PCR for CDH1, VIM, SNAIL, ZEB1 and NFATc1before and after treatment with siRNAs against NFATc1. (C) Western blot for CDH1, VIM, ZEB1, SNAIL in H1975 cells. β-actin is used as loading control. Left column represents the control siRNA, second and third columns represent two independent siRNAs against NFATc1. *, p < = 0.01 using two tailed t-test.*

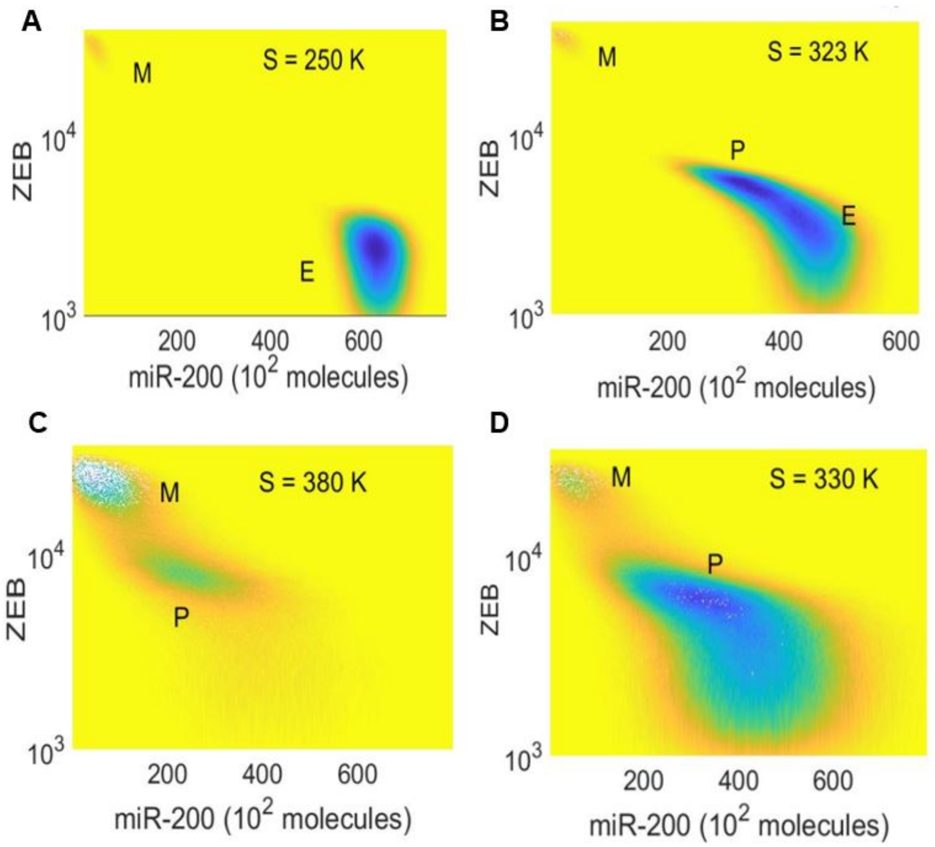


***FigS3: Potential landscape for NFATc-EMT coupled circuit at varying SNAIL levels***

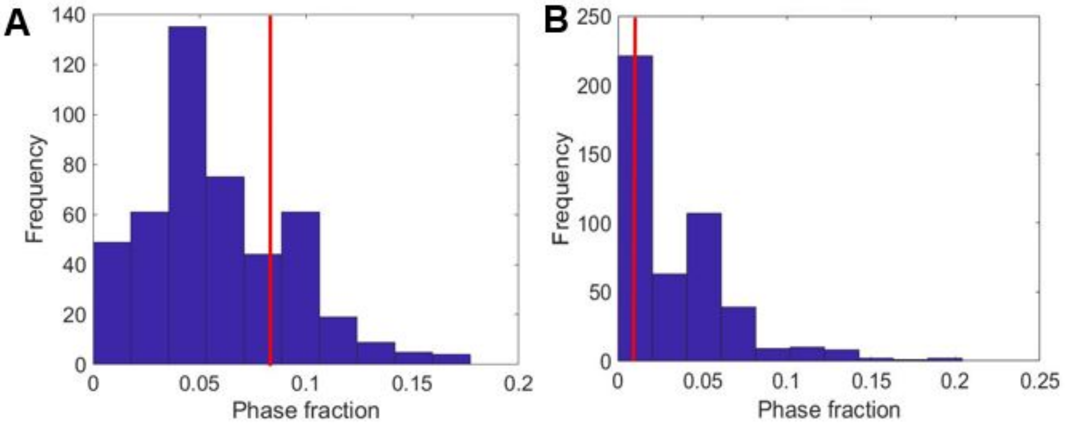


***FigS4: Phase analysis of randomized networks.*** *A) Frequency distribution of {E, H} phase fraction for 561 randomized networks. The red line denotes the phase fraction of {E, H} phase in the wild type (WT) NFATc-EMT coupled network. (B) Frequency distribution of {H, M} phase fraction for 561 randomized networks. The red line denotes the phase fraction of {H, M} phase in the wild type (WT)*

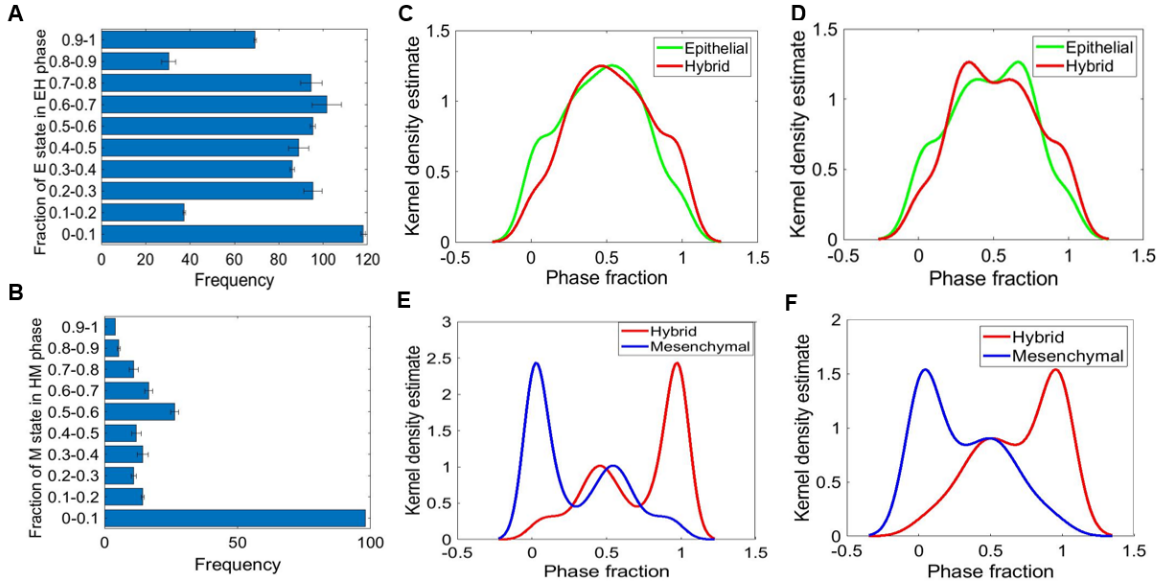


**FigS5: Relative stability analysis.** *A) Frequency distribution of E state in {E,H} phase. B) Frequency distribution of M state in {H,M} phase. C-D) Density plots showing the distribution of E and H state in the {E,H} phase for two parameter sets obtained from independent RACIPE replicates. E-F) Same as C-D but for H and M states in the {H, M} phase.*

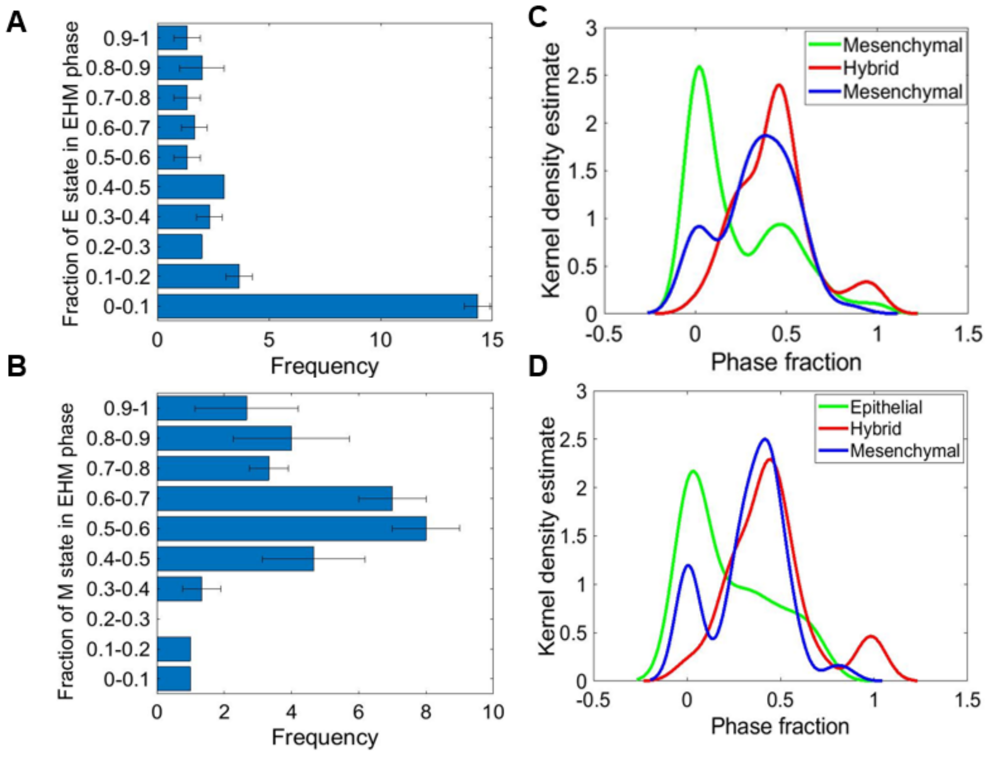


***FigS6: Relative stability analysis**. A) Frequency distribution of E state in the {E,H,M} phase. (B) Frequency distribution of M state in the {E,H,M} phase. C-D) Density plot showing the distribution of E, H and M state in the {E,H,M} phase for two parameter sets obtained from two independent RACIPE replicates/runs.*

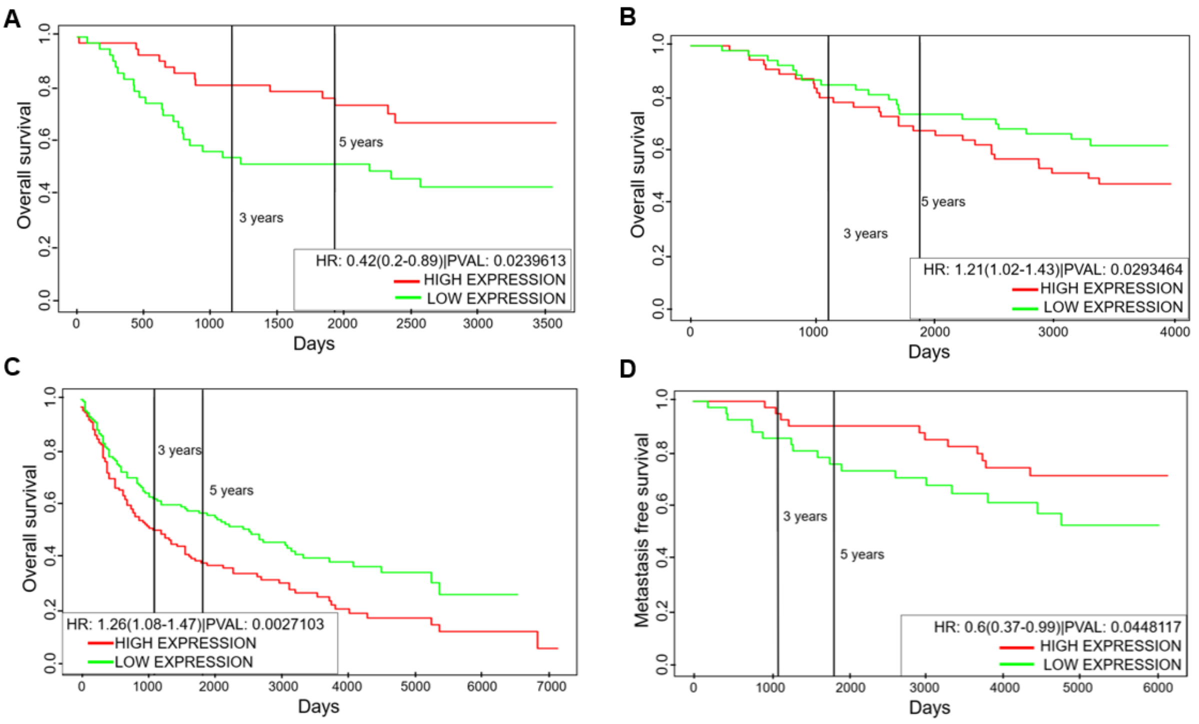


**FigS7:** ***Clinical outcome of NFATc1 levels is tissue dependent***. *(A) Overall survival for GSE14814 (Lung cancer sample). B-C) Overall survival for GSE19536 (Breast cancer samples) and GSE30219 (Lung cancer samples). D) Metastasis free survival for GSE6532 (breast cancer samples). Group of patients with high NFATc1 is denoted by red curve; those with low NFATc1 is depicted by green curve. All cohorts are divided based on median levels.*

**References:**


1. Lu M, Jolly MK, Levine H, Onuchic JN, Ben-Jacob E. MicroRNA-based regulation of epithelial-hybrid-mesenchymal fate determination. *Proc Natl Acad Sci U S A* (2013) **110**:18174–9. doi:10.1073/pnas.1318192110

2. Mooney SM, Jolly MK, Levine H, Kulkarni P. Phenotypic plasticity in prostate cancer: role of intrinsically disordered proteins. *Asian J Androl* (2016) **18**:704–10. doi:10.4103/1008-682X.183570

3. Gould R, Bassen DM, Chakrabarti A, Varner JD, Butcher J. Population Heterogeneity in the Epithelial to Mesenchymal Transition Is Controlled by NFAT and Phosphorylated Sp1. *PLoS Comput Biol* (2016) **12**:e005251. doi:10.1371/journal.pcbi.1005251

4. Oikawa T, Nakamura A, Onishi N, Yamada T, Matsuo K, Saya H. Molecular and Cellular Pathobiology Acquired Expression of NFATc1 Downregulates E-Cadherin and Promotes Cancer Cell Invasion. (2013) doi:10.1158/0008-5472.CAN-13-0274

5. Serfling E, Chuvpilo S, Liu J, Höfer T, Palmetshofer A. NFATc1 autoregulation: a crucial step for cell-fate determination. *Trends Immunol* (2006) **27**:461–469. doi:10.1016/j.it.2006.08.005
